# The digestive system of a cricket pulverizes polyethylene microplastics

**DOI:** 10.1101/2023.05.23.541961

**Authors:** Marshall W. Ritchie, Jennifer F. Provencher, Jane E. Allison, Matthew J. Muzzatti, Heath A. MacMillan

## Abstract

Microplastics (MPs; <5 mm) are a growing concern and a poorly understood threat to biota. Despite a recent spike in research on MPs, most of this work has focused on marine systems, and less is known about interactions between terrestrial organisms. We used a generalist insect (a cricket; *Gryllodes sigillatus*) to examine whether individuals would ingest and physically degrade MPs in their food. We fed crickets a range of concentrations (0, 2.5, 5, and 10% w/w) of fluorescent MPs mixed into a standard diet and dissected the gut regions to isolate the MPs within. Comparing plastic content and fragment size within gut regions, we sought to identify whether and where crickets can fragment ingested MP particles. Given the digestive tract morphology of this species, we expected that the crickets would both ingest and egest the MPs. We also predicted that the MPs would be fragmented into smaller pieces during this digestive process. We found that *G. sigillatus* egested much smaller pieces than they ingested (likely into the nanoplastic range), and this fragmentation occurs early in the digestive process of this insect. These findings suggest that generalist insects can act as agents of plastic transformation in their environment if/when encountering MPs.

## Introduction

Plastic is a widely used and ever-growing resource worldwide, with production increasing substantially since mass production began in the mid-20th century (1). While today, plastic is mass-produced at 320 million tons annually (2,3), recycling efforts to deal with waste plastics are limited throughout the world (4). An estimated 80% of plastic is not recycled, and waste plastics frequently enter natural environments (2,5–7). Despite the attention aquatic environments receive, not all plastic pollution ends up in marine ecosystems. Since the 1950s, about 55% of all plastics have been sent to landfills and other terrestrial areas. An estimated 4900 megatons of plastics have been directed to terrestrial systems thus far (6,7) suggesting that terrestrial environments play a role in the fate of plastic pollution.

When plastics are introduced to the natural environment, they begin to degrade. This degradation can be caused by abiotic and biotic factors such as weather, tidal cycles, sunlight, and animal interactions (5,8). Through these processes, plastics can physically break down into microplastics (MPs), plastics smaller than 5 mm in size (5), with poorly understood impacts (9). While there has been a rapid increase in research on MPs in recent years, most of this work has examined MP impacts on marine animals (10). The potential for terrestrial animals to interact with MPs has largely been ignored (11). This is a huge gap in our understanding of how organisms may interact with plastics given that MP contaminated soil can reach 6.7% w/w in industrialized areas (12).

One of the first invertebrate terrestrial MP studies by Huerta Lwanga et al., (2016) shed light on the little-known effects MPs have on terrestrial animals as they examined how contaminated soil impacts a terrestrial species. They found earthworms (*Lumbricus terrestris*) in MP-contaminated soil had reduced growth rates with increasing MP concentrations in their environment. The potential for such effects is not limited to earthworms; a study by Lu et al., (2020) more recently found that 95% of animals representing 20 wild terrestrial species (including insects and crustaceans) contained MPs. These species had ingested a range of MP shapes, from spheres to fibres. However, research testing how MPs can affect the physiology and fitness of terrestrial species is limited, as is research on the passage of MPs through the digestive tract of terrestrial invertebrates and their subsequent fate in the environment.

Many insects, like crickets, are prey for larger animals (including vertebrates) and typically ingest small animal debris or plant material on the soil surface (15). *Gryllodes sigillatus* is a terrestrial cricket species with worldwide distribution that has been used as a model organism to study microplastic ingestion (Thomas, 1985; Otte, 2006; Fudlosid et al., 2022). In a laboratory environment, this species has been previously shown to readily ingest and excrete microplastic beads and fibres throughout development when the plastics were mixed into their feed (19). Although plastic ingestion had no obvious negative effects on these insects in this initial study, ground-dwelling species may play important roles in plastic transformation in natural environments.

The cricket gut is divided into three major functional regions, the foregut, midgut, and hindgut, which contribute to the breakdown of food via mechanical and/or chemical degradation (20–22). The proventriculus is a muscular grinding organ located at the posterior end of the foregut, important in the mechanical breakdown of hard foods (20,21). Plastics are generally resistant to chemical degradation, so in addition to mastication, grinding organs like the proventriculus may strongly contribute to any breakdown of plastics that might occur in the insect gut (23).

Here, we examined the passage and transformation of MPs moving through the gut of a generalist insect to address three hypotheses relating to the fate of the plastics. First, we hypothesized that crickets (*G. sigillatus*) would ingest small MPs indiscriminately and predicted that larger doses of plastic mixed into their feed would result in larger quantities of plastic found within the entire gut. Second, we hypothesized that the gut of a generalist omnivore insect could break plastics down into smaller-sized fragments (physical transformation), thereby allowing such insects to serve as agents of plastic transformation in their environment. Third, we hypothesized that such transformation of plastics, should it occur, would happen mainly in the foregut through the grinding actions of the proventriculus.

## Materials and Methods

### Cricket rearing

All crickets were reared under either long-term or short-term plastic exposure using methods adapted from Ritchie et al. (2022). Tropical house crickets (*Gryllodes sigillatus*) were reared from eggs acquired from Entomo Farms (Norwood, ON). Eggs were held in an incubator at 32°C and 40% RH with a 14:10 L:D light cycle. Once the eggs hatched, nymphs (1^st^ instar cricket) were randomly assigned to bins following the methods outlined in Ritchie et al. (2022). Juvenile crickets were left for one week in these bins to grow and develop beyond this fragile early stage before being assigned to treatment groups.

To allow for long-term plastic feeding (exposure to plastic throughout development), crickets were assigned to new bins at one week of age, where they were exposed to either the control diet or the same base diet but containing 2.5%, 5%, or 10% w/w of microplastic (MP). The MPs mixed with the food were low-density Fluorescent Blue Polyethylene Microspheres ∼100 μm (Cospheric, Fluorescent Blue Polyethylene Microspheres 1.13 g/cc 90-106 μm). In each bin, 40 juvenile crickets of unknown sex were placed with the food. Crickets were provided unlimited water, food (respective to their diets), and egg cartons for shelter throughout development. Rearing of the crickets was finished when the crickets were between 7 and 8 weeks of age. Each week the bins were cleaned, and survival was quantified.

To examine the effects of short-term plastic feeding on adults, crickets were reared as in the long-term plastic feeding experiment and were assigned to new bins immediately after the first week of development. Upon this transfer, 40 juvenile crickets were counted and fed the control diet until they reached the experimental age of 6 to 7 weeks. Crickets were provided food and water *ad libitum* and cardboard egg cartons for shelter throughout development. Adult crickets were randomly selected to receive the same dietary concentrations of plastic as in the long-term developmental exposure experiment. They were then sorted by sex and randomly transferred into new clean bins with a maximum of 20 crickets of mixed sex with their respective MP diets for 3, 6, or 9 days of plastic feeding. At the end of their respective short-term feeding, they had reached the age of 7 to 8 weeks. Survival was recorded at the end of feeding to test whether longer plastic exposure times led to greater cricket mortality.

### Cricket gut dissection

All cricket dissections were carried out using the methods described in Ritchie et al. (2022). When crickets from both the long- and short-term exposure experiments finished their allotted plastic exposure time, they were at least seven weeks old and dissected within one week. Each individual was weighed using a microbalance (Sartorius ME5 model) and anesthetized with CO_2_ (15s); they were placed in a second petri dish of saline (NaCl, CaCl_2_ (dihydrate), NaHCO_3_, Glucose, KCl, recipe as in Whitaker et al., 2014) after being washed (saline, DI H2O) to remove any MPs that may be on the outside of the exoskeleton. The cricket was then carefully dissected in the saline under a dissecting microscope. The cricket gut was then removed and placed on a dry petri dish, where it was segmented into the foregut, midgut, and hindgut (anatomy and sections, as shown in Figure 1). The segmenting was done on a dish outside of the saline to ensure all MPs within the gut were not washed away. The proventriculus of the crickets was pulled apart to allow the plastic to be removed in the KOH digestion steps. Each region was weighed using a microbalance (Sartorius ME5 model) and frozen at -20 °C.

**Figure 1:**
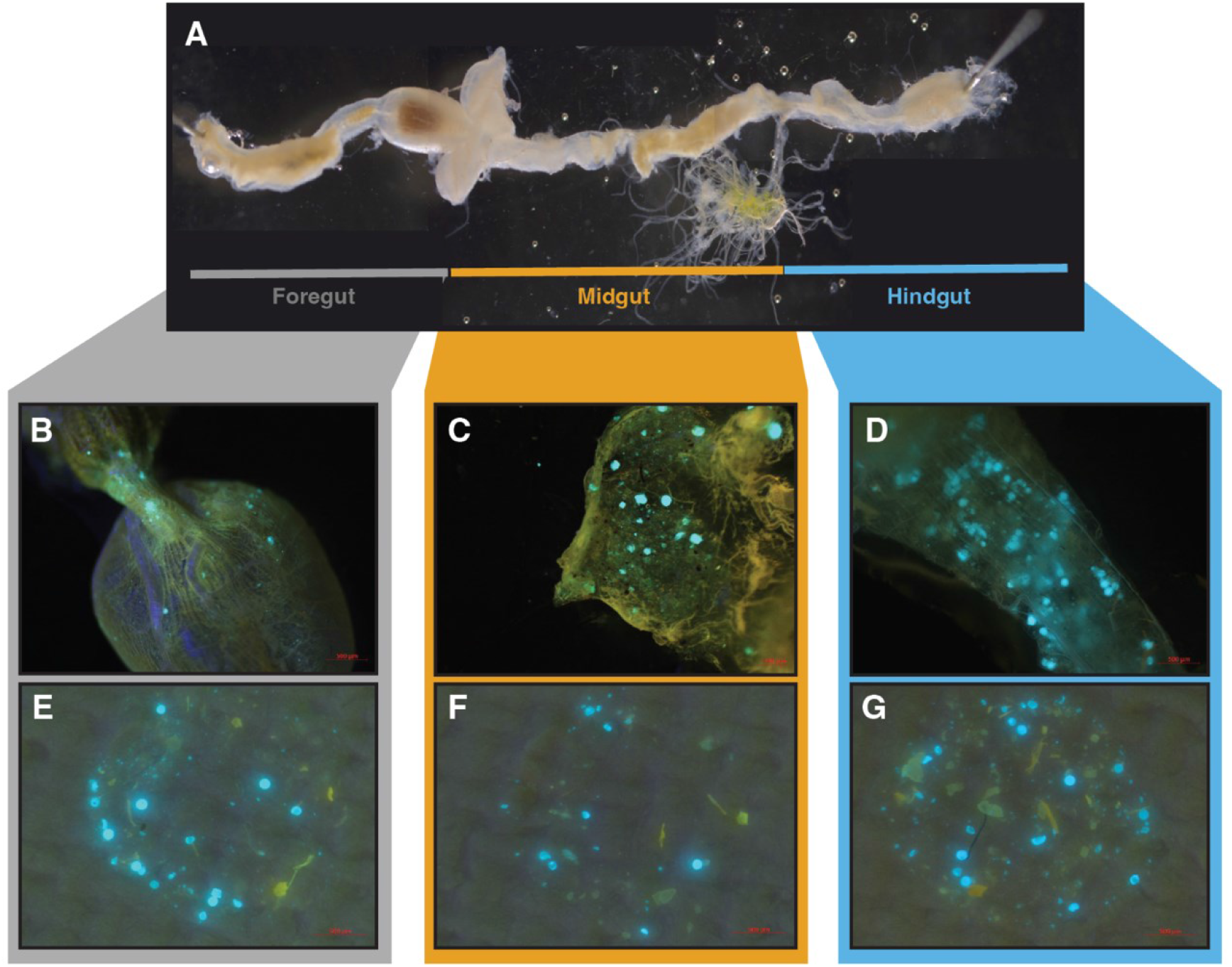
Composite photo of the complete digestive tract of an adult female cricket, *Gryllodes sigillatus* (without plastics) (A) and representative images of gut regions containing plastics before (B-D) and after (E-G) the sample digestion process (representative images from different samples). Images of filters with isolated fluorescent plastics (E-G) show the amount of plastic obtained from applying a 5 μL subsample (from a 150-250 μL sample) of a digested gut region to each filter.

### Microplastic isolation

The samples from each group were removed from the freezer and thawed (26). Once thawed, 10% KOH was added to the samples (200 μL for the foregut, 250 μL for the midgut and plastic controls, and 150 μL for the hindgut), and they were left for two days at 60°C to digest (27). KOH was provided to raw beads to unsure there was no breakdown caused by the digestion method. After the two days, the samples were briefly vortexed to ensure a homogeneous solution, and 5 μL samples of each sample were pipetted onto a 1 μm pore size glass fibre filter (Sigma, APFB04700) on top of a vacuum pump assembly (Sigma, Z290408-1EA). Pressure was applied to the system using a hand pump (Fisher, S12932) attached to the vacuum pump assembly. After filtering, the filter paper, along with the MPs trapped on it, was moved to a petri dish and photographed under a dissection microscope (Zeiss, Oberkochen, Stemi 508 with an axiocam 105 colour) fitted with a 450 nm long-pass emission filter and 400-415 nm excitation light source (Nightsea LLC, Lexington, United States, Stereo Microscope Fluorescence Adapter). After this, the filter paper with the MPs was inverted on a clean petri dish, and the smallest particle visible was identified visually and photographed using an inverted microscope (Zeiss, Vert.A1 with an axiocam 202 mono). The inverted microscope was fitted with a LED-Module 385 nm and a FL Filter Set 49 for emission and excitation of the MPs beads. An exposure time limit of max 75 ms and 150 ms was set for the dissecting and inverted microscopes, respectively, as early trials revealed that images that had higher values could falsely inflate particle sizes. These thresholds are specific to the microscopes used.

### Image analysis

Image analysis was carried out following the methods described in Ritchie et al. (2022). Images of fluorescent plastic particles were analyzed using ImageJ (FIJI version, 2.3.051; java, 1.8.0_172[64-bit]). Particles in the samples with areas larger than 20000 μm^2^ were removed as this was just over double the expected largest bead area. The 20000 μm^2^ cut-off removed 608 out of 20410 captured samples (∼3%) from the analysis. An informal analysis of these images confirmed that these images contained particles that were too close together and thus resulted in an implausible particle area. Of the 570 filter images, 49 (∼9%) could also not be processed as there was too much plastic to accurately determine the area of individual pieces or plastics that were too small and could not be seen using this camera. No fluorescence signal was detected in any of the control samples.

Due to the dissection scope limitations in detecting the smallest particles (24), we opted to use a fluorescence-equipped inverted microscope that allowed us to view the smallest particles visible in each sample. This additional step was done using only samples from female crickets. Images from the inverted microscope were analyzed similarly to those from the dissecting microscope, except that RBG stacking and Z project function was not used, as the original photos were black and white. In replacement for these steps, the image was changed to an 8-bit image, then auto threshold using the steps defined above. See Ritchie et al. (2022) for a fully detailed method for plastic detection.

### Data analysis

The total plastic visible area (μm^2^) within a sample was calculated by multiplying the subsample area by the dilution factor and dividing by the gut region’s mass. All graphing and statistical analysis were completed in R version 4.1.2 (28). To understand which of the predictor variables of time, gut region, sex, and plastic concentration most strongly influenced the amount of plastic found within the gut and how these predictor variables interacted, we used linear mixed effects models and AIC for model selection. The fixed effects were time, gut region, sex, and concentration, and the random effect was the ID of the cricket. We found that the best fitting model excluded the amount of time crickets had fed on the plastic as a fixed effect. We had a large sample size in this experiment, so while this model provided the best fit, almost all possible interactions among the variables were statistically significant. Thus, we further explored relationships among the variables and the strength of effects to understand the most biologically significant interactions represented in the figures.

## Results

To characterize how passage of MPs through the gut of a ground-dwelling invertebrate change the plastics, we dissected and isolated the gut’s three major anatomical and functional regions (foregut, midgut, and hindgut; Figure 1A) after confirming that plastic ingestion did not contribute to cricket mortality during rearing (Table S1). Through visual inspection of the gut, we observed the presence of plastic within each of the gut regions of the crickets (Figure 1B, C, D), indicating that the crickets were feeding on the plastic-augmented diets and not avoiding ingesting the plastics. Using the filtering methods described in Ritchie et al. (2022), we isolated MPs in a subsample of digested material from gut regions using vacuum filtration with a 1 μm pore size (Figure 1E, F, G).

After filtering, we used an automated imaging (ImageJ) and counting process to characterize the fluorescent particles in all three gut regions of the 190 crickets exposed to a plastic diet (we identified no particles in any control samples) (24). The total area of plastic within each sample was then corrected by dilution factor and tissue mass to get the total area of plastic from each individual relative to gut region mass (Figure 2). Using a model selection approach (AIC), we found that total plastic particle area relative to gut mass (an index of the total amount of plastic within the gut region, relative to the mass of the region) was significantly impacted by sex, plastic dose, and gut region with no interactions, and with no effect of time spent feeding on the diet (Table S2). In this model, we found that the gut region explained the largest amount of variation in total area of plastic relative to cricket mass (F_2,319_ = 43.03, p < 0.0001), with plastic dose (F_2,184_ = 20.0, p < 0.0001), and sex (F_1,183_ =12.97, p = 0.0004), each explaining less variation than the gut region (in that order; supplemental Table 3).

**Figure 2:**
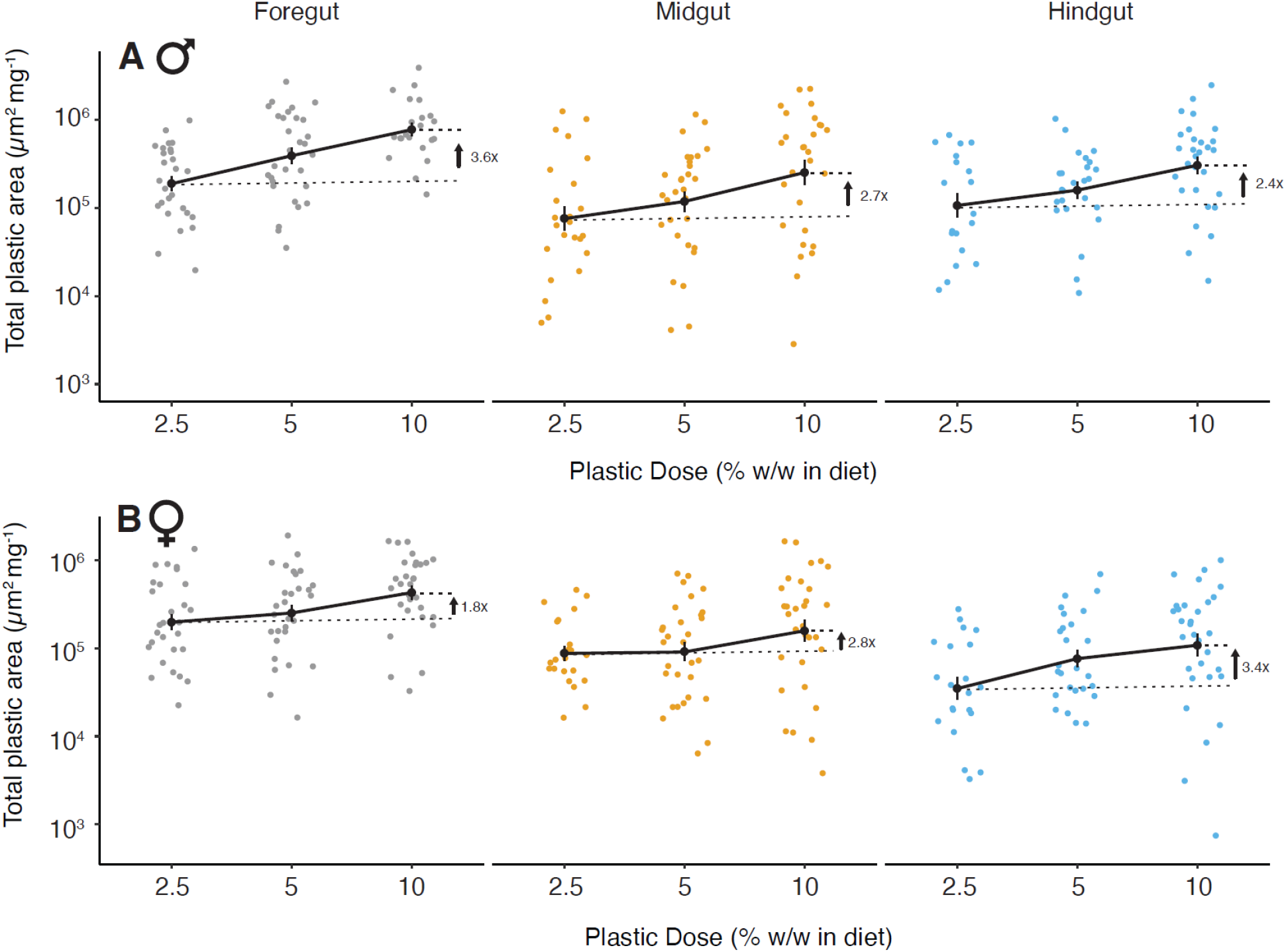
Total plastic area (μm^2^/mg) found inside each gut region (foregut, midgut, and hindgut) relative to tissue mass in male and female crickets. As the concentration of plastic in the diet increased, the total quantity of plastic within gut regions similarly increased for both sexes. Individual crickets’ total estimated area of plastic is shown by which concentration of plastic feed in each was given. The coloured dots represent the total area of plastic calculated for individual gut regions, and the black dots represent the mean total estimated area of plastic ± SE of all the individuals.

As the concentration of plastic increased 4-fold in the diet, the mean total area of plastic found in the guts of both male and female crickets increased by approximately 1.8- and 3.6-fold (Figure 2). The amount of plastic contained within each gut region decreased along the digestive tract, with the foregut containing the greatest amount of total plastic (Figure 2). These trends likely drive the significant effects of gut region and plastic concentration in the diet seen in both males and females (Supplemental Table 3). Males had less total plastic within their foreguts compared to females, while the hindguts of females had less than males.

In addition to estimating the quantity of plastic in the gut, we also quantified the sizes of particles found in each gut region. The vast majority of observed particles were smaller than the initial size of the beads added to the diet, indicating that the crickets were fragmenting most of the ingested plastic particles. Overall, there was an approximately 10-fold change in the mean size of plastic particles found within the gut of crickets compared to control (un-ingested) beads (Figure 3). The amount of time crickets fed on plastics did not influence the amount of breakdown within the gut (based on model AIC), so time spent feeding on plastic was excluded from the best-fitting model (Supplemental Table 4). As with the total amount of plastic contained within the gut, we found that the gut region explained the largest amount of variance in particle size (F_2,16577_ = 136.06, p < 0.0001; supplemental Table 5), indicating that additional plastic breakdown occurs as particles progress through the gut lumen. The foregut was the region that contained the largest fragments, while the midgut and hindgut contained similarly sized fragments overall. There were also significant interactions in the effects of gut region, dose, and sex on particle size (F_4,16898_ =19.94, p < 0.0001; Supplemental Table 5).

**Figure 3:**
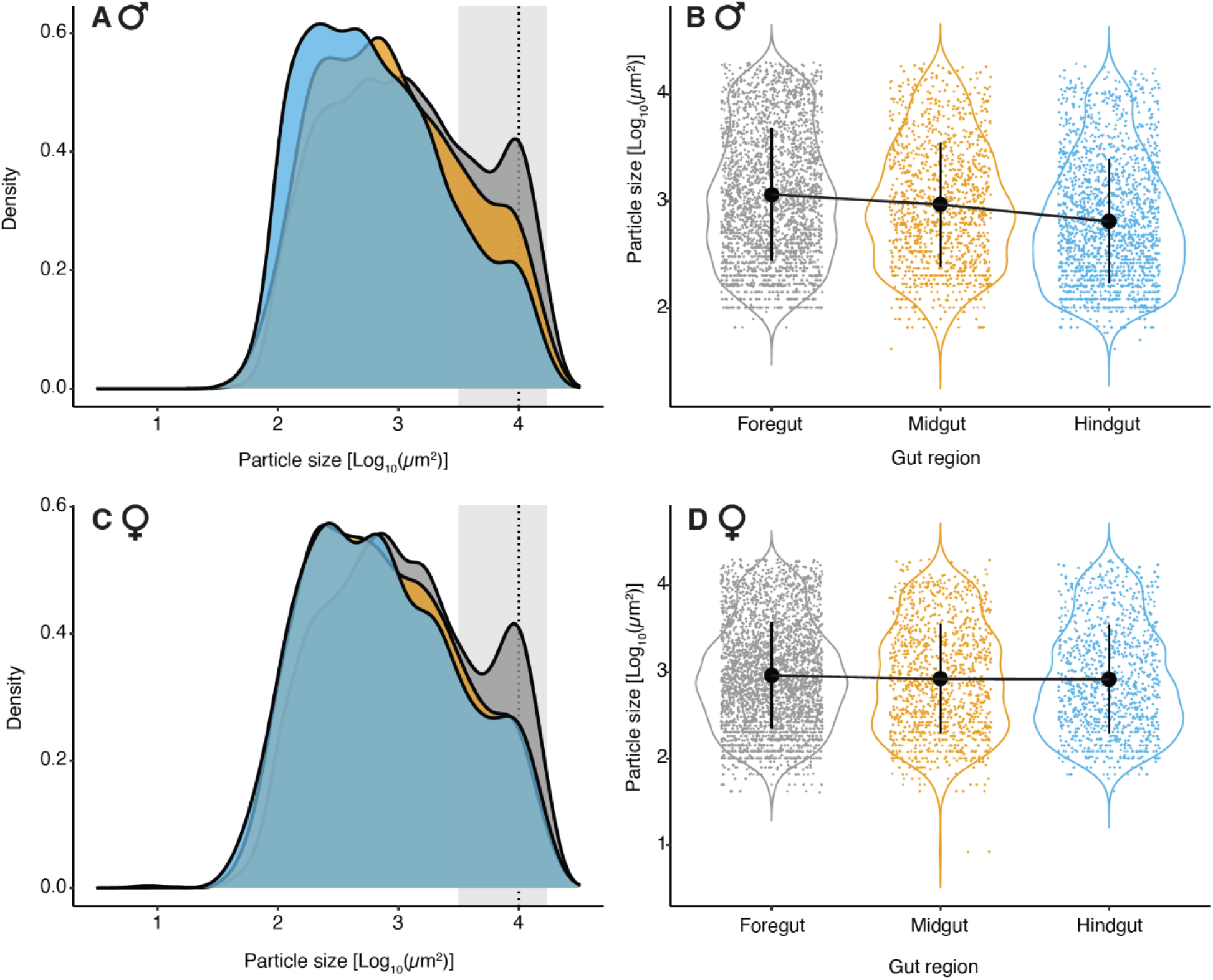
Male crickets were able to break down MPs to smaller sizes as they moved through the digestive tract, while the majority of breakdown occurred before or in the foregut of females. Area density plots of the gut regions for male and female crickets (A and C). Individual pieces of plastic contained within each gut region, using only 10.00% plastic, fed results (B and D). Grey represents the foregut, yellow is the midgut, and blue is the hindgut. The dashed line shows the mean KOH treated bead area (10000 μm^2^), and the shaded area is the standard deviation (± 6782).

Given that gut region explained the majority of variation of the particle size and interacted with sex (F = 8.32, p < 0.0001; supplemental Table 5), we compared the location and degree of breakdown in males and females separately. We found that males tended to have fewer whole beads within all the gut regions than females (Figure 3A, C). Females tended to have a larger number of whole beads in the foregut, and an equal level of plastic breakdown was observed in the midgut and hindgut. Compared to the females, males tended to have broken down the MPs more once they had reached the hindgut (Figures 3A and C). The males, on average, had smaller particles as plastic was moving down the gut regions and this was most clear in crickets that fed on a diet that was 10% w/w plastic (Figures 3B and D). Males on average decreased the area in the mean (black dot) through the gut regions (Figure 3B). Females, however, did not have the same trend, and the mean particle size was consistent through the gut regions (Figure 3D). These trends were also seen in the 2.50% and 5.00% fed crickets but were most apparent in the 10.00% group. This explains our significant three-way interaction between gut region, plastic dose, and sex (Supplemental Table 5). A notable drop-off in particles detected that are less than 42 μm^2^ in area is caused by our predetermined detection limit (equivalent to an area of 5 pixels) when using a dissecting microscope. Polyethylene beads that were exposed to the same KOH treatment were measured within one standard deviation, as highlighted in the shaded gray area (Figure 3A and C).

To examine whether crickets break down MP beads to sizes smaller than those we could detect with a dissecting microscope (42 μm^2^), we analyzed a subset of the same samples using an inverted microscope with greater magnification (Figure 4A). Again, the smallest particles that were detected were frequently at or close to the limit of detection of our methods. In this case, the area of the smallest MPs identified was 1.47 μm^2^ equivalent to the predetermined threshold of 5 pixels, as previously mentioned (Figure 4B). This area also equates to a particle diameter of approximately 1.35 μm, which is very close to the pore size of the filters used to extract our samples (1 μm). Particles smaller than this size would be nanoplastics and would not be captured by filtering.

**Figure 4:**
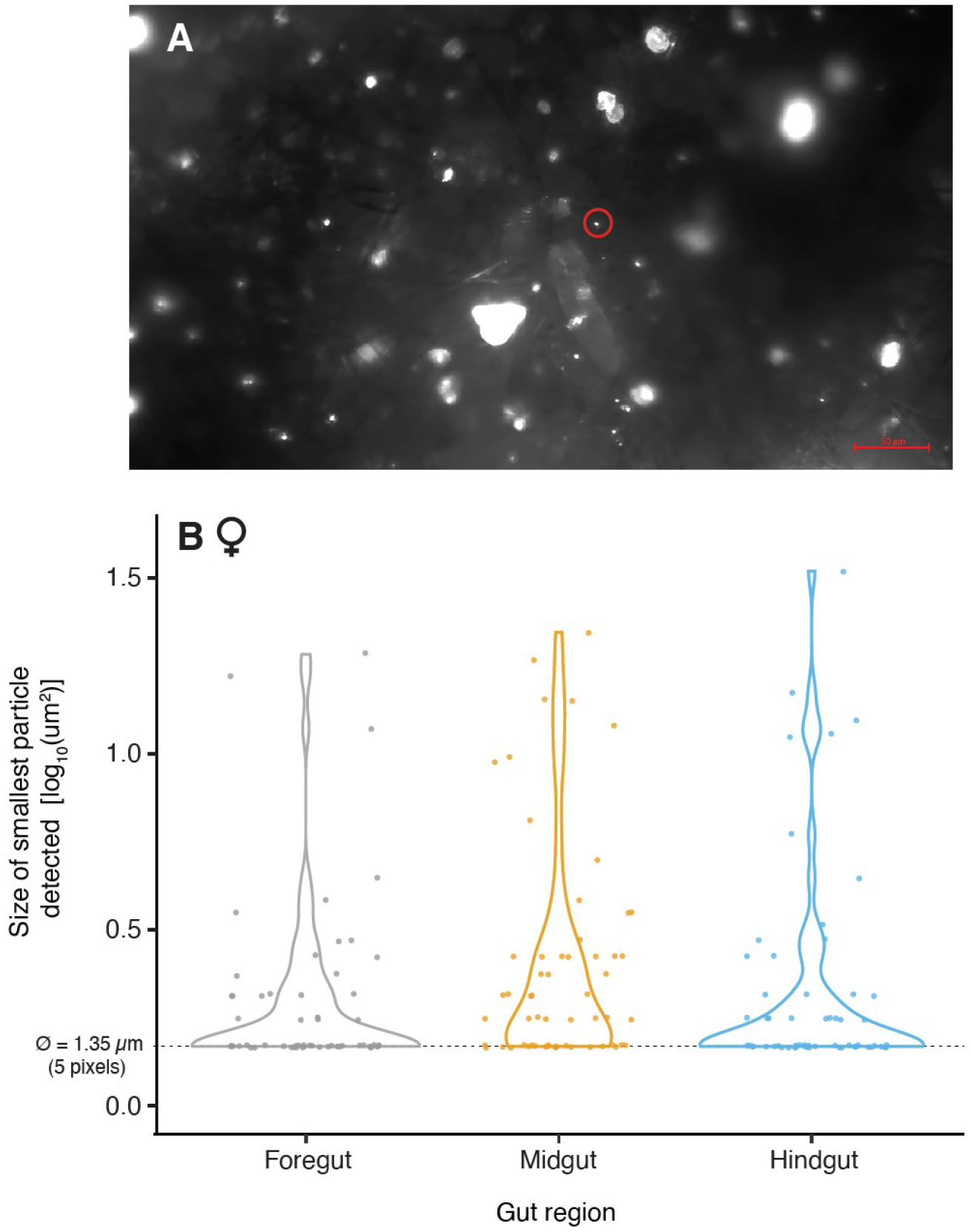
The smallest particles identified from photos of gut contents taken using an inverted microscope. A representative image from the inverted microscope (A). Female cricket’s smallest particle size is seen on the filter in all the gut regions (B). Violin plots show the distribution of the smallest particle sizes among the images, and points represent are the smallest fluorescent particle identified from each sample. The dashed line highlights where the smallest particles are being detected at the area of 1.43 μm^2^ (diameter of 1.35 μm), which is set by our image analysis threshold of 5 pixels. The red circle in panel A highlights one of these small particles.

## Discussion

Exposure of terrestrial species to MPs will become more common as plastic waste continues to pollute terrestrial systems. When plastics enter natural environments, they can undergo a degradation process from abiotic factors, such as hydrolysis and light exposure (5,11), leading to MPs and possibly even nanoplastics (NPs; plastics smaller than 0.1 μm in size (29)). Our results reveal that biotic factors, such as ingestion by a terrestrial insect (*G. sigillatus*), are also capable of causing the mechanical breakdown of MPs into smaller MPs, and likely even NPs, and do not chemically alter the polymers. The change in chemical composition of the plastics were tested with plastics manually extracted from the frass and was found to have not been chemically changed (Table S6).

Previously, Fudlosid et al., (2022) quantified the effects of polyethylene ingestion on growth rates and survival in the same cricket species and found that *G. sigillatus* growth was not affected by up to 10% of their diet being untreated polyethylene beads. Our findings of the present study support these results; we saw no obvious effect of plastic feeding on cricket survival (Supplemental Table 1).

Although plastic feeding had no effect on cricket growth and survival, passage through the gut of a cricket had dramatic effects on the plastics. Several factors impacted how the plastics were altered on their passage through the cricket gut. As the concentration of plastic in the diet increased, so did the total amount of plastic found within the gut regions (Figure 2; Supplemental Table 3). This implies that crickets can and do ingest very high concentrations of plastics in their diets in a lab environment, and the amount of plastics is influenced by the level they are exposed to. We did not measure the amount of food consumed by the crickets as we fed them *ad libitum* and thus cannot discern from our experiments if the crickets would have avoided plastic food if given a choice of food without plastic. It is important to note, however, that a four-fold plastic increase was not found when comparing the smallest and largest concentrations. This suggests that the crickets could avoid the plastics at higher concentrations or separate the food at smaller concentrations (Figure 2). It is also possible that the filter size chosen resulted in us missing particles smaller than 1 μm, such that the total quantity of plastic in the final sample was skewed. Further research should explore whether insects avoid plastic in their diets using controlled diet choice experiments.

Our second hypothesis was that crickets could break plastics into smaller fragments, thereby acting as agents of plastic transformation in their environments. We found clear evidence of the extensive breakdown of polyethylene particles throughout each of the gut regions of the crickets, with the largest fragments of beads residing in the foregut (Figure 3A, C). On average, plastics within the gut that we were able to quantify with a dissecting microscope were approximately 10-fold smaller in size compared to the size of the MPs mixed in their feed (Figure 3B, D). Using the more powerful inverted microscope to measure the smallest particles, we found particles that were nearly a 1000-fold smaller than the original plastic (Figure 4), which suggests that the cricket gut is capable of converting microplastics down to the nanoplastic scale. Taken together, these data suggest that crickets can serve as significant agents of plastic transformation within their environment and the can result as few as three days. We suggest that the willingness of crickets to ingest plastics in the laboratory, their capacity to transform plastics to smaller sizes, and their ability to survive plastic exposure all are likely to increase the probability of transfer of these MPs up through trophic levels (e.g., through predation), or down (e.g., from detritivores ingesting insect frass). While the concentrations that these crickets were fed likely do not represent the levels of plastic contamination most common in the environment, even the smallest concentration in our study (2.5%) yields the same conclusion, and is within the range of levels of plastic contamination already reported in soils in industrial areas (0.3-6.7% w/w (12)). Notably, there are also reports of some terrestrial environments reaching extremely high levels of plastic contamination (>1100 particles per gram dry weight), therefore even the highest levels of exposure may be relevant in some highly polluted soils (30).

It should be noted that we quantified many particles at or near our limit of quantification. This suggests that the crickets are mechanically breaking down the plastic particles to even smaller sizes that reach the Nanoplastic (NP) size range. Nanoplastics are notoriously difficult to isolate and quantify (31), and while we chose a filter mesh size (1 μm) that ultimately controlled for the lower extreme of plastic size. We could only isolate down to this threshold, and how many NPs are generated within the gut relative to the larger particles quantified here remains unclear. Given the increased concern about NPs in biota, and the potential effects on a range of processes, future work should focus on NPs in terrestrial systems, including how insects may be contributing to NP transfer in food webs.

We expected that plastic breakdown within the gut could be caused mainly by the strong muscle-grinding organ, the proventriculus (Figure 1). If the proventriculus caused the breakdown, we expected to see a substantial reduction in the size of plastic particles between the foregut (anterior to and including the proventriculus) and the midgut (posterior to the proventriculus). Instead, a large proportion of the breakdown appears to occur before or within the esophagus and crop (before the proventriculus), suggesting that the cricket mouthparts also contribute significantly to the breakdown observed. In Antarctic krill, the most likely location of fragmentation was reported to be physical breakdown by the mandibles, the stomach, and the gastric mill (a grinding organ similar to the proventriculus) (32). Biofragmentation has also recently been demonstrated in mealworms, and involved both physical and chemical degradation by the mouthparts, microbiome, and digestive enzymes (33). Given this, and our own results (Figure 3), we therefore argue that the proventriculus contributes to, but is not solely responsible for, the breakdown of MPs within the insect gut (Figure 3A, C). Insect diets and gut anatomy are diverse, and different gut regions likely can play different roles in MP degradation in different taxa. Recently, Helmberger et al., (2022) reported that crickets could fragment polystyrene, but they faced methodological limitations in characterizing the extent or location of this breakdown within the crickets. These similarities between the locations of the breakdown of MPs by crickets and other species (Antarctic krill and mealworms, (32,33)) provide further evidence that hard structures along the gut, such as the mandibles or proventriculus, may be the primary contributors to physical fragmentation of plastics.

We did not observe evidence of plastic accumulating in the gut over time (Supplemental Table 1), nor evidence of plastics being retained within a particular region of the gut. These observations could suggest that crickets pass plastic beads and fragments through their digestive tract and excrete most or all of the breakdown products back into their environment. By contrast, mealworms were found to retain plastics in their digestive tract for more than two days after feeding (33), implying plastics may slow clearance in some species, but not others.

Crickets readily ingest polyethylene microplastic beads and grind them down to produce at least 1000-fold smaller particles very early in the digestion process. These small plastics do not appear to be blocking or building up in the gut. If crickets similarly ingest and breakdown plastics in nature, we argue that this can lead to trophic transfer of even smaller plastic particles up to predators of crickets. Crickets may also facilitate the uptake of micro- and nanoplastics by much smaller animals via the excretion of these fragmented plastics. When plastic exits the cricket digestive system, it is very likely that some pieces are at the nanoplastic scale, and there is limited information available thus far on the effects of NPs on biota. A large variety of insects have chewing mandibles and a grinding proventriculus that aids in the breakdown of hard foods common to their diet. We, therefore, argue that the ability to breakdown plastic is unlikely to be unique to *G. sigillatus*, and that insects may serve as an important pathway for plastic transformation in contaminated environments.

## Acknowledgements

The authors would like to thank Mads Anderson for the support in discussions about the filtering process and imaging with the inverted microscope. The authors would also like to thank Hannah Anderson for assisting with cricket care. We would also like to thank Entomo Farms for supplying cricket eggs during the project’s development.

## Author Contributions

All authors conceived the study and designed the experiments. MR carried out the experiments. MR and HAM curated and analyzed the data and created visualizations. HAM and JFP provided resources and supervision. MR drafted the manuscript, and all authors edited the manuscript.

## Data availability

All data is provided as a supplementary file for review, and the same file will be included as supplementary material should the manuscript be accepted for publication.

