## Supplemental tables for "The digestive system of a cricket pulverizes polyethylene microplastics"

**Supplementary Material**

**The digestive system of a cricket pulverizes polyethylene microplastic beads**

Ritchie et al.

**Table 1:** Survival of crickets fed different concentrations of plastics, separated by sex and experiment duration. The numerator shows the amount surviving after treatment, and the denominator shows the total number of crickets originally assigned to each treatment group.

| Concentration (w/w) % | 3 days | | 6 days | | 9 days | | Long term | |
|---|---|---|---|---|---|---|---|---|
| | *Female* | *Male* | *Female* | *Male* | *Female* | *Male* | *Female* | *Male* |
| 0 (control diet) | 85.7% 12/14 | 94.4% 17/18 | 93.3% 14/15 | 100% 15/15 | 100% 14/14 | 100% 16/16 | 52.5% (21/40) 10 | 11 |
| 2.5 | 93.3% 14/15 | 100% 18/18 | 100% 15/15 | 100% 15/15 | 100% 17/17 | 93.3% 14/15 | 65% (26/40) 11 | 15 |
| 5 | 93.8% 15/16 | 100% 18/18 | 92.3% 13/14 | 100% 17/17 | 94.1% 16/17 | 100% 17/17 | 55% (22/40) 8 | 14 |
| 10 | 100% 15/15 | 100% 18/18 | 92.9% 13/14 | 87.5% 14/16 | 100% 17/17 | 100% 17/17 | 65% (26/40) 10 | 16 |

*Note: Long-term survival percentage is the total number of crickets that survived; sex was unknown until crickets reached adulthood. Forty newly hatched juveniles were placed in each bin for the long-term condition. Short-term survival tracking started at plastic exposure and ended at dissection.*

**Table 2**: Output from mixed-effects models on total plastic area relative to gut mass. In all cases, data were $log_{10}$-transformed before analysis, individual cricket ID was treated as a random effect, and parameters were estimated by restricted maximum-likelihood (REML). K = the number of parameters in the model. AIC = Akaike information criterion. ΔAIC = difference in model AIC relative to the best fitting model (in bold). Res.LL = restricted log-likelihood. For the model parameters, Gut = foregut, midgut, or hindgut, Plastic = concentration of plastic in feed excluding controls, and Time = length of the feeding period in days.

| Model parameters | K | AIC | ΔAIC | Res.LL |
|---|---|---|---|---|
| **Sex+Plastic+Gut** | **8** | **782.7** | **0.0** | **-383.3** |
| Time+Sex+Plastic+Gut | 11 | 792.0 | 9.4 | -385.0 |
| Sex*Gut | 8 | 800.9 | 18.3 | -392.5 |
| Sex+Gut | 6 | 802.2 | 19.6 | -395.1 |
| Gut | 5 | 805.1 | 22.5 | -397.6 |
| Sex*Plastic*Gut | 20 | 816.1 | 33.4 | -388.0 |
| Plastic | 5 | 880.4 | 97.8 | -435.2 |
| Time | 4 | 889.9 | 107.3 | -441.0 |
| Sex | 6 | 903.2 | 120.5 | -445.6 |
| Time*Sex*Plastic*Gut | 74 | 922.8 | 140.1 | -387.4 |

**Table 3:** Output from linear mixed effects model of best fit describing the effects of the gut region, plastic concentration in the feed (Plastic dose), and sex on total plastic area corrected by gut region mass. Cricket ID was included as a random effect. Parameters in bold were significant predictors of the total plastic contained within the gut.

| Model parameters | Sum Sq | Mean Sq | NumDF | DenDF | F value | P value |
|---|---|---|---|---|---|---|
| **Gut region** | **1E+13** | **5E+12** | **2** | **319** | **43.03** | **<0.0001** |
| **Plastic dose** | **4.65E+12** | **2.32E+12** | **2** | **184** | **20.00** | **<0.0001** |
| **Sex** | **1.51E+12** | **1.51E+12** | **1** | **183** | **12.97** | **0.0004** |

**Table 4:** Output from mixed-effects models on particle size. In all cases, data were $\log_{10}$-transformed before analysis, individual cricket ID was treated as a random effect, and parameters were estimated by restricted maximum-likelihood (REML). K = the number of parameters in the model. AIC = Akaike information criterion. ΔAIC = difference in model AIC relative to the best fitting model (in bold). Res.LL = restricted log-likelihood. For the model parameters, Gut region = foregut, midgut, or hindgut, Dose = concentration of plastic in feed, and Time = length of the feeding period.

| Model parameters | K | AIC | ΔAIC | Res.LL |
|---|---|---|---|---|
| **Sex*Dose*Gut region** | **20** | **35839.9** | **0** | **-17900** |
| Gut region | 5 | 35869.7 | 29.8 | -17929.9 |
| Sex+Gut region | 6 | 35875.7 | 35.8 | -17931.9 |
| Sex+Dose+Gut region | 8 | 35880.2 | 40.3 | -17932.1 |
| Sex*Gut region | 8 | 35880.4 | 40.5 | -17932.2 |
| Time*Sex*Dose*Gut region | 74 | 35892.1 | 52.2 | -17872.1 |
| Time+Sex+Dose+Gut region | 11 | 35898.8 | 58.8 | -17938.4 |
| Dose | 5 | 36226.8 | 386.9 | -18108.4 |
| Sex | 4 | 36228.8 | 388.9 | -18110.4 |
| Time | 6 | 36240.2 | 400.3 | -18114.1 |

**Table 5:** Output from linear mixed effects model of best fit describing the effects of the gut region, plastic concentration (Dose), and sex on particle size. Cricket ID was included as a random effect. Parameters in bold were significant predictors of particle size within the gut.

| Model parameters | Sum Sq | Mean Sq | NumDF | DenDF | F value | P value |
|---|---|---|---|---|---|---|
| **Gut region** | **3.42E+09** | **1.71E+09** | **2** | **16,577** | **136.06** | **<0.0001** |
| **Dose** | **9.93E+07** | **4.97E+07** | **2** | **164** | **3.95** | **0.021** |
| Sex | 8.42E+06 | 8.42E+06 | 1 | 166 | 0.67 | 0.414 |
| **Gut region*Dose** | **2.89E+08** | **7.23E+07** | **4** | **16,898** | **5.75** | **<0.0001** |
| **Gut region*Sex** | **2.09E+08** | **1.05E+08** | **2** | **16,577** | **8.32** | **<0.0001** |
| Dose*Sex | 9.55E+05 | 4.78E+05 | 2 | 164 | 0.04 | 0.963 |
| **Gut region*Dose*Sex** | **1.00E+09** | **2.51E+08** | **4** | **16,898** | **19.94** | **<0.0001** |

**Table 6:** FTIR results of untreated plastic from the bottle and plastic was that manual extracted from frass. There appears to be no difference between bottle and frass plastic matching.

| Compound/chemical | Positive matches from untreated plastic sample (/5) | Positive matches from frass sample (/5) | Untreated plastic match (%; mean +/- SD) | Frass plastic match (%; mean +/- SD |
|---|---|---|---|---|
| Docosanol | 5 | 5 | 93.24 +/- 0.33 | 91.4 +/- 1.94 |
| Polyethylene (low density) | 5 | 5 | 92.17 +/- 0.70 | 90.4 +/- 2.15 |
| *N-Octadecyltrimethylammonium chloride* | 5 | 5 | 90.24 +/- 0.54 | 89.04 +/- 1.73 |
| Polyethylene | 4 | 5 | 89.32 +/- 0.38 | 88.32 +/-1.12 |
| Triacontane | 4 | 4 | 89.46 +/-1.01 | 88.01 +/- 2.08 |
| *N-Hexadecyltrimethylammonium chloride* | 1 | 1 | 87.99 +/- NA | 86.67 +/- NA |

FTIR analysis, results, and discussion

Fourier transform infrared spectroscopy (FTIR) using a Nicolet iN10 Infrared Microscope (Fisher, Massachusetts) was used for material identification. We used this approach to confirm the chemical identity of virgin plastic from the bottle from Cospheric, as well as plastic that was manually extracted from frass pellets of crickets fed 10% (w/w) plastic throughout their development. The mounting process of each plastic was done by labelling the underside of a glass slide into a grid to allow for easy placement and was then cleaned with filtered 1 μm distilled water. This was to ensure all plastics placements were known and all possible contaminates were removed with the water. After the glass slide was dry, a thin layer of skin tac (Skin-Tac Liquid Adhesive Barrier) was applied on the top slide and under the dissection microscope, each plastic was placed using forceps. All ten pieces of plastic were mounted on one slide. The top 5 matches of each plastic were found to have been matched to one of the following libraries of HR Hummel Polymer and Additives, Aldrich Condensed Phase Sample Library, and HR Nicolet Sampler Library.

An FTIR spectrometer was used to identify plastics supplied by the manufacturer as well as those found in frass. We found no chemical differences in the polyethylene fragments (Supplemental Table 6). The top five chemical matches for each of five samples were all very similar and had high confidence matches. They indicated MPs showed no change in chemical composition from the crickets passing them through their entire digestive tract.

Our hypothesis was that plastics passing through the cricket gut would remain chemically unaltered by their passage. Using FTIR, we found no chemical differences between virgin plastic from the manufacturer's bottle and the plastic that was physically removed from frass pellets (Supplemental Table 6). This result suggests that the breakdown of the MPs we observed was likely caused primarily by the cricket gut's physical interaction with the plastic. There has been some investigation into the ability of terrestrial animals, such as earthworms and snails, to chemically break down ingested MPs (1,2). In both these studies, the plastic breakdown was caused by the gut microbiome of these species rather than physical alterations. The crickets may similarly contain enzymes or microbes capable of plastic breakdown that impact the surface of the plastics within the gut, but we do not think this is the leading cause of the breakdown seen.
